# Heterogeneity in local density allows a positive evolutionary relationship between self-fertilisation and dispersal

**DOI:** 10.1101/203042

**Authors:** James Rodger, Pietro Landi, Cang Hui

## Abstract

Theoretical work predicts that dispersal and self-fertilisation (selfing) should always be negatively correlated and the Good Coloniser Syndrome (GCS) of high dispersal and selfing should not occur when both traits are free to evolve. This contradicts positive relationships between selfing and dispersal in empirical data. Critically, previous work assumes density of adults is spatially and temporally homogeneous, so selfing results in homogeneity in propagule production and competition, which eliminates the benefit of dispersal for escaping from local resource competition. We investigate the joint evolution of dispersal and selfing in a demographically structured metapopulation model where local density varies due to stochastic extinction-recolonisation dynamics. Increasing local extinction rate reduces local density across the metapopulation, which favours high selfing to mitigate mate limitation, but increases heterogeneity in density, which favours high dispersal for escape from competition. Together, these effects produce a positive relationship between selfing and dispersal, and evolution of the GCS. Nevertheless, the relationship between selfing and dispersal is context-dependent, as varying dispersal cost yields a negative relationship. Our results imply that if spatiotemporal heterogeneity in environmental suitability increases towards the range edge, the GCS may evolve there, favouring further range expansion (Cf. Baker’s Law).

## 1. Introduction

Both self-fertilisation (selfing) and dispersal affect gene flow and colonisation success, which in turn have strong effects on geographical distributions, biological invasions and speciation (Busch 2011, Massol and Cheptou 2011b, Hargreaves and Eckert 2014, Pannell 2015, Hui and Richardson, 2017). Therefore, elucidating constraints that enforce joint evolution of selfing and dispersal, and lead to the emergence of mating-dispersal syndromes, would have broad implications (Cheptou and Massol 2009).

Baker’s Law (Baker 1955, Stebbins 1957) states that species which can self-fertilise should be better colonisers because selfing assures reproduction when mate or pollinator availability limits outcross reproduction in the new environment. Based on this, it can be expected that highly dispersive, frequently colonising species should benefit from selfing, leading to evolution of the Good Coloniser Syndrome (GCS) of high dispersal and high selfing (argument developed by Cheptou and Massol 2009). However, quantitative theoretical investigations accounting for the effect of selfing rate on dispersal rate evolution and vice versa predict a strictly negative relationship between these traits, precluding evolution of the GCS (Cheptou and Massol 2009, Massol and Cheptou 2011a, Sun and Cheptou 2012). This contrasts to empirical studies showing positive relationships of selfing ability with dispersal ability (Darling et al. 2008, De Waal et al. 2014). Moreover, the theoretical studies only predict syndromes of dispersal with cross-fertilisation (outcrossing) or selfing with no dispersal. This is inconsistent with larger native and invaded range sizes in species with greater selfing ability, given that ranges must be occupied by dispersal (Grossenbacher et al. 2015; van Kleunen and Johnson, 2007). Notwithstanding that theory focuses on rates and empirical work on abilities for selfing and dispersal (Pannell et al. 2015), these opposite patterns warrant further investigation. As certain model assumptions of the theoretical studies are highly restrictive with respect to dispersal evolution, models with different assumptions may better accommodate empirical results.

Selection on dispersal depends on the relative advantages of offspring staying versus leaving their site of origin (Clobert et al. 2004, Ronce 2007). In the metapopulation framework, which has been widely used for theoretical studies of dispersal, dispersal rate is defined as the probability of a propagule (the dispersing stage, e.g. a seed) emigrating from its patch. In general, spatiotemporal heterogeneity selects positively on dispersal rates (Levin et al. 1984, McPeek and Holt 1992) because the patches in which many of the propagules are produced in one generation are not the best patches for them to complete their lives and reproduce in the next generation (Clobert et al. 2004, Ronce 2007, Kubisch et al. 2014). For example, under heterogeneity in density-dependent competition, dispersal can be favoured because it reduces competition (Comins 1980, Parvinen 2006). This benefit is usually referred to as “escape from competition”, even though dispersal reduces competition for both dispersed and nondispersed propagules. Dispersal may also be favoured for avoidance of kin competition (Hamilton and May 1977) and inbreeding (Motro 1991, Perrin and Mazalov 1999), especially if the population is genetically structured (has spatial genetic heterogeneity). Negative selection on dispersal comes from costs, such as the risk of failing to reach destination habitat after leaving the habitat of origin (cost of dispersal: Hamilton and May 1977). In addition, where small populations suffer mate limitation (i.e., under an Allee effect), individuals that reach vacant habitat may reproduce poorly if only small numbers of propagules arrive, selecting against dispersal (Robinet and Liebhold 2009). This last result comes from a study on insects (gypsy moths) which are always unisexual and therefore cannot evolve selfing. However, in a system where selfing is possible, selection could favour increased selfing instead of reduced dispersal, potentially giving rise to the Good Coloniser Syndrome (GCS) of high dispersal and high selfing.

While most studies of selfing focus on continuous populations (e.g., Lande and Schemske 1985, Charlesworth and Charlesworth 1987, Cheptou and Fenster 2004, Morgan and Wilson 2005), investigating the implications of colonisation for selfing evolution requires a spatially structured setting, such as a metapopulation. Generally, three main factors govern selection on selfing (Lloyd 1992, Barrett 2010, Eckert et al. 2006, Karron et al. 2012). Reproductive assurance is the benefit of selfing in mitigating mate or pollinator limitation. In metapopulations, this should favour evolution of selfing when empty patches are colonised by small numbers of propagules (Pannell and Barrett 1998, Dornier et al. 2008), as mate limitation frequently occurs in small and sparse populations (Leimu et al., 2006, Gascoigne et al. 2009). This is an example of the Allee effect, defined generally as a reduction in performance due to low abundance (positively density-dependent performance, Stephens et al. 1999). Cheptou and Massol (2009), on the other hand, explore evolution of selfing in response to pollinator limitation unrelated to density in a metapopulation. Selfing is also promoted by the transmission advantage: that selfers can both self and outcross, and so pass on more copies of their genes to the next generation than strict outcrossers (Fisher 1941). Inbreeding depression, the poorer performance of offspring from inbreeding than outcrossing, opposes evolution of selfing (Darwin 1876, Charlesworth and Charlesworth 1987, Husband and Schemske 1996) but can itself evolve to lower levels when self-fertilisation rates remain high for several generations (Lande and Schemske 1985, Barrett and Charlesworth 1991, Crnokrak and Barrett 2002).

Despite the rich theoretical literatures on the evolution of selfing and dispersal individually (see reviews by Clobert et al. 2004, Ronce 2007, Barrett 2010, Karron et al. 2012), few models have considered the evolution of both traits simultaneously (Ravigne et al. 2006, Cheptou and Massol 2009). The previously studied model of Cheptou and Massol (2009), which we refer to as the heterogeneity in pollinators (HP) model, represents a metapopulation where presence versus absence of pollinators fluctuates in habit patches but all patches have identical density of adult plants. Increasing the rate of pollinator failure (absence of pollinators) selects for higher selfing. However, as selfing rate increases, heterogeneity in seed production and consequently local resource competition is reduced (i.e., the number of seeds produced by patches with versus without pollinators becomes more similar), which selects for lower dispersal. Hence the HP model predicts that selfing and dispersal should be negatively related and the GCS should not occur (Cheptou and Massol 2009, Massol and Cheptou 2011a, Sun and Cheptou 2012). However, it has not always been sufficiently emphasised (e.g., Massol and Cheptou 2011a, Sun and Cheptou 2012, Auld and de Casas 2013) that the effect of selfing on selection of dispersal in the HP model, which results in the negative relationship, is specific to situations where selfing mitigates pollinator limitation (i.e. plants only) and local density is identical across patches (Busch 2011, Massol and Cheptou 2011b).

In this study, we investigate the joint evolution of selfing and dispersal in a model where heterogeneity in density arises from stochastic local extinction followed by recolonisation and population growth. Selfing is selected to assure reproduction under mate limitation, which affects both animals and plants. Previous investigations in the same general modelling framework, but with only one trait evolving, show that positive selection on dispersal arises from heterogeneity in local density (Parvinen 2006), and positive selection on selfing arises from the presence of recently recolonised patches with low density (Dornier et al. 2008). This suggests that a positive evolutionary relationship between selfing and dispersal, and hence evolution of the GCS, may occur in this framework when the two traits evolve simultaneously.

## 2. Model and methods

### 2.1. Metapopulation model

We adapted and extended the metapopulation models of Parvinen (2006) for evolution of dispersal and Dornier et al. (2008) for evolution of selfing to allow evolution of both traits simultaneously. We refer to our model as the heterogeneity in density (HD) model, as local density is heterogeneous due to stochastic local extinction and recolonisation. The HD model considers a hermaphrodite species with an annual life history in a metapopulation of an infinite number of identical patches occupied by local populations. Time is discrete and events in each time step (year) are as follows.

- Individuals produce female gametes (ovules for plants or ova for animals), a fraction of which are self-fertilised (prior selfing, Lloyd 1992). Due to inbreeding depression, a fraction of self-fertilised ovules/ova fail to produce propagules (for example seeds, fertilised eggs or actively dispersing young).
- The ovules which are not self-fertilised are available for cross fertilisation. The fraction of these ovules which is actually cross-fertilised (and produces propagules) is positively related to local density (i.e., there is an Allee effect due to mate limitation).
- Of all propagules produced, a fraction disperses (emigrates) and the remainder stay in the local population.
- A fraction of dispersed propagules is lost (the cost of dispersal), and successfully dispersed propagules are evenly distributed among all patches, following the island model of dispersal of Hamilton and May, 1977.
- After emigration and immigration, carrying capacity of patches imposes competition (establishment from propagule to adult is thus negatively density dependent).
- Finally, a fraction of local populations suffer extinction due to environmental stochasticity. Individuals in the remaining populations go on to produce propagules in the following year.

Events thus happen each year in the following order: reproduction, emigration, immigration, competition, and local extinction. The metapopulation has an age distribution of local populations *p_*τ*_ = e*(l — *e*)^*τ*^, where *e* is local population extinction rate and *τ* the age of these populations, with *τ* = 0 for newly extinct populations (Ronce et al. 2001). In a local population, per capita reproduction (*λ*) is described by

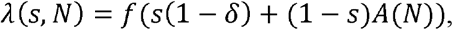

where *f* is the number of ovules produced per individual (fertility), selfing rate *s* the fraction of these which is self-fertilised, and *δ* the fraction of self-fertilised ovules dying due to inbreeding depression (Cheptou 2004). (1 – *s*)*A*(*N*) ovules are cross fertilised, where *A(N)* = N/(*a* + *N*) due to the Allee effect in mate limitation (Cheptou 2004), where *N* is the local population density and parameter *a* the local population density at which half of the ovules/ova available for outcrossing are actually outcrossed. The density of propagules in each local population following emigration and immigration is:

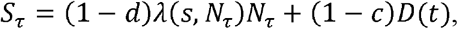

where dispersal rate *d* is the fraction of propagules dispersing (emigrating) and *c* is the fraction of these lost during dispersal (cost of dispersal). As all habitat patches are assumed to be physically identical, local population density is equivalent to size. Here we refer to density to be consistent with the terminology of competition and Allee effects as density dependent processes.

Metapopulation viability is assessed from the dispersal pool *D*(*t*), which gives a measure of size of the metapopulation as whole, analogous to mean local density (Parvinen 2006, Dornier et al. 2008). If the dispersal pool falls to zero, the metapopulation becomes extinct. The dispersal pool depends on the age distribution of local populations and is calculated as

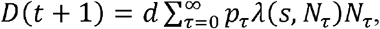

where *t* represents the succession of years in the metapopulation (Dornier et al. 2008). Due to competition, survival of these propagules to adulthood in the following year is density dependent. Per-capita survival is thus *C*(*S*) = 1/(1 + *bS*), with 1/*b* being the carrying capacity of each local population (Cheptou 2004). The number of adult plants *N* in populations is thus described by,

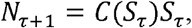

with *N_*τ* = 0_* = 0. Thus, reproduction is governed by the Allee effect in mate limitation and survival is governed by competition.

Estimation of fitness in this model is simplified in that it ignores the transmission advantage of self-fertilisation (Fisher 1941). In this respect we underestimate selection on self-fertilisation, so our assessment of conditions favouring evolution of self-fertilisation, and hence the GCS, is conservative. Nevertheless, a model incorporating the automatic transmission advantage indicates that this does not qualitatively affect our results (Supplementary Materials).

### 2.2. Methods

We study the eco-evolutionary HD model to assess the effects of environmental and demographic parameters on evolution of dispersal rate *d* and selfing rate *s,* and on ecological extinction or viability. The parameters investigated are local extinction rate *e,* intensity of the Allee effect in outcrossing success *a,* intensity of competition *b,* cost of dispersal *c*, inbreeding depression *δ* and fertility *f*. Each simulation begins with (1) a set of parameter values; (2) starting values for dispersal rate and selfing rate *(d, s);* and (3) an initial size of the dispersal pool *D.* In the initial part of the simulation, the metapopulation is allowed to reach demographic equilibrium, i.e., when *D* remains constant between generations. The size of the equilibrium dispersal pool is then evaluated to check whether the metapopulation remains viable or has gone extinct. As in the Dornier (2008), our model reveals three ecological scenarios, depending on the set of parameter values and initial selfing and dispersal rates. (1) The metapopulation becomes extinct, regardless of starting density (Unconditional Extinction). (2) The metapopulation is viable, regardless of starting density (Unconditional Viability). (3) The outcome depends on the starting density, with extinction occurring if the total number of individuals in the metapopulation at the starting point is below a threshold and viable otherwise (Conditional Viability) (Dornier et al. 2008). The Conditional Viability Scenario indicates an emergent Allee effect at the level of the metapopulation (Dornier et al. 2008). In our simulations, we distinguish between these three scenarios for each combination of parameter values and trait values (*d, s*) investigated by starting simulations from very low and very high initial values of *D* (Dornier et al. 2008). If both cases lead to extinction, this indicates Unconditional Extinction; if both lead to viability, this indicates Unconditional Viability; and if starting at low *D* leads to extinction but starting at high *D* leads to viability, this indicates Conditional Viability.

We investigate evolution of selfing and dispersal rates following standard adaptive dynamics procedure (Dieckmann and Law 1996, Metz et al. 1996, Dercole and Rinaldi 2008). Rare mutant individuals characterised by traits (*d', s'*) are introduced into the metapopulation with resident traits (*d, s*) at its demographic equilibrium. If the mutants have higher fitness than the residents, the mutant traits become fixed, and thus become the resident traits for the next round of selection. This process is repeated until an evolutionary singularity is reached, i.e., where the selection gradient is neutral (slope of the fitness landscape = 0). Evolutionary singularities occur either at fitness maxima, where evolution stops, indicating an Evolutionarily Stable Strategy (ESS) or at fitness minima, when selection is disruptive, leading to evolutionary branching (Geritz et al. 1997, 1998, Della Rossa et al. 2015, Dercole et al. 2016). In our model, evolutionary singularities were always fitness maxima. Thus, the system does not display polymorphism through evolutionary branching. This is because with monotonic density-dependent functions, the age-size distribution is also monotonic, which prevents the time-heterogeneity needed for evolutionary branching (Parvinen 2006). Evolutionary suicide is also never observed in our model, consistent with Parvinen (2006), since evolution of *s* and *d* always proceeds away from the Unconditional Extinction region. We also never find alternative evolutionary singularities but always a single global ESS. We therefore discuss how the ESS changes in different environmental and demographic conditions. A detailed explanation of the computation of the relative reproductive fitness (invasion fitness) of the mutant is provided in Appendix A.

As *s* and *d* both range from 0 to 1, they can be visualised on a plane divided into four quadrants separated by the lines for *d* = 0.5 and *s* = 0.5. We treat these quadrants as different evolutionary syndromes: Ld-Ls (low dispersal-low selfing), Ld-Hs (low dispersal high selfing), Hd-Ls (high dispersal-low selfing), and Hd-Hs (high dispersal-high selfing), which we interpret as the GCS. We choose 0.5 as a cut off because it is the midpoint of the potential range of both traits in our model. This is reasonable for selfing, which almost always evolves to the boundaries (0, 1) in our model and has a similar but continuous bimodal distribution in nature. In the context of our model, this is also reasonable for dispersal rate, but in natural systems, where this trait may be constrained in a smaller but unknown range (for instance through constraints on evolution of dispersal structures), dispersing-selfing syndromes are probably better evaluated in relative terms (see Darling et al. 2008, De Waal et al. 2014).

To assess the likelihood of finding the various ecological scenarios and evolutionary syndromes in our model, we performed simulations with randomly sampled parameter values and initial trait values of *d* = 0.5 and *s* = 0.5. First, for a sample of 100 simulations, we recorded the ecological scenario at demographic equilibrium. Second, for 100 simulations that did not begin in the Unconditional Extinction scenario, we recorded in which quadrant of (*d, s*) space the ESS occurred. To investigate the effects of variation in parameters on the joint evolution of selfing and dispersal rates and hence the location of the ESS, we performed a sensitivity analysis with the ESS located at *d* = 0.5 and *s* = 0.5. While the HD model does not allow analytical solutions relating parameters to selection gradients and ESS, our definition of fitness (Section 2.1) allows an intuitive interpretation, and the approximated analysis of selection (Appendix B), which does have analytical solutions, provides some insights into the full HD model.

## 3. Results

### 3.1 Simulations with randomly chosen parameter values

Simulations with randomly chosen parameter values give evolutionary endpoints in all four quadrants of (*d, s*) space. These endpoints were always ESS. With respect to the ecological scenarios, 5% of simulations started in the Unconditional Viability scenario, 70% in the Conditional Viability scenario, and 25% in the Unconditional Extinction scenario. Of simulations not starting in the Unconditional Extinction scenario, 50% converged to the GCS (Hd-Hs), 32% to Ld-Hs, 10% to Hd-Ls, and only 8% to Ld-Ls. The ESS for selfing rate was nearly always 1 or 0 (complete selfing or complete outcrossing) but the ESS for dispersal never approached 1 or 0 (complete dispersal or complete non-dispersal). Although randomly chosen parameter values did not produce ESS of intermediate selfing rates (0 < *s* < 1), this is possible, but only under a very narrow range of parameters (Fig. S1).

We show selected simulations to illustrate effects of parameter values on evolutionary and ecological outcomes (Figs 1, 2). In these examples, the GCS is favoured by low cost of dispersal *c* with high Allee effect parameter *a* (Fig. 1b) and by high local extinction rate *e* with low inbreeding depression *δ* (Fig. 2b). In the supplementary materials we show that omitting the transmission advantage of selfing does not qualitatively affect these results (Fig. S2). The area of (*d, s*) space falling into the Unconditional Extinction Scenario becomes larger with decreasing propagule production and increasing mortality: i.e., as Allee effect parameter *a,* competition parameter *b,* inbreeding depression *δ,* cost of dispersal *c*, and local extinction rate *e* increase and as fertility *f* decreases (Figs 1, 2).

**Fig. 1.**
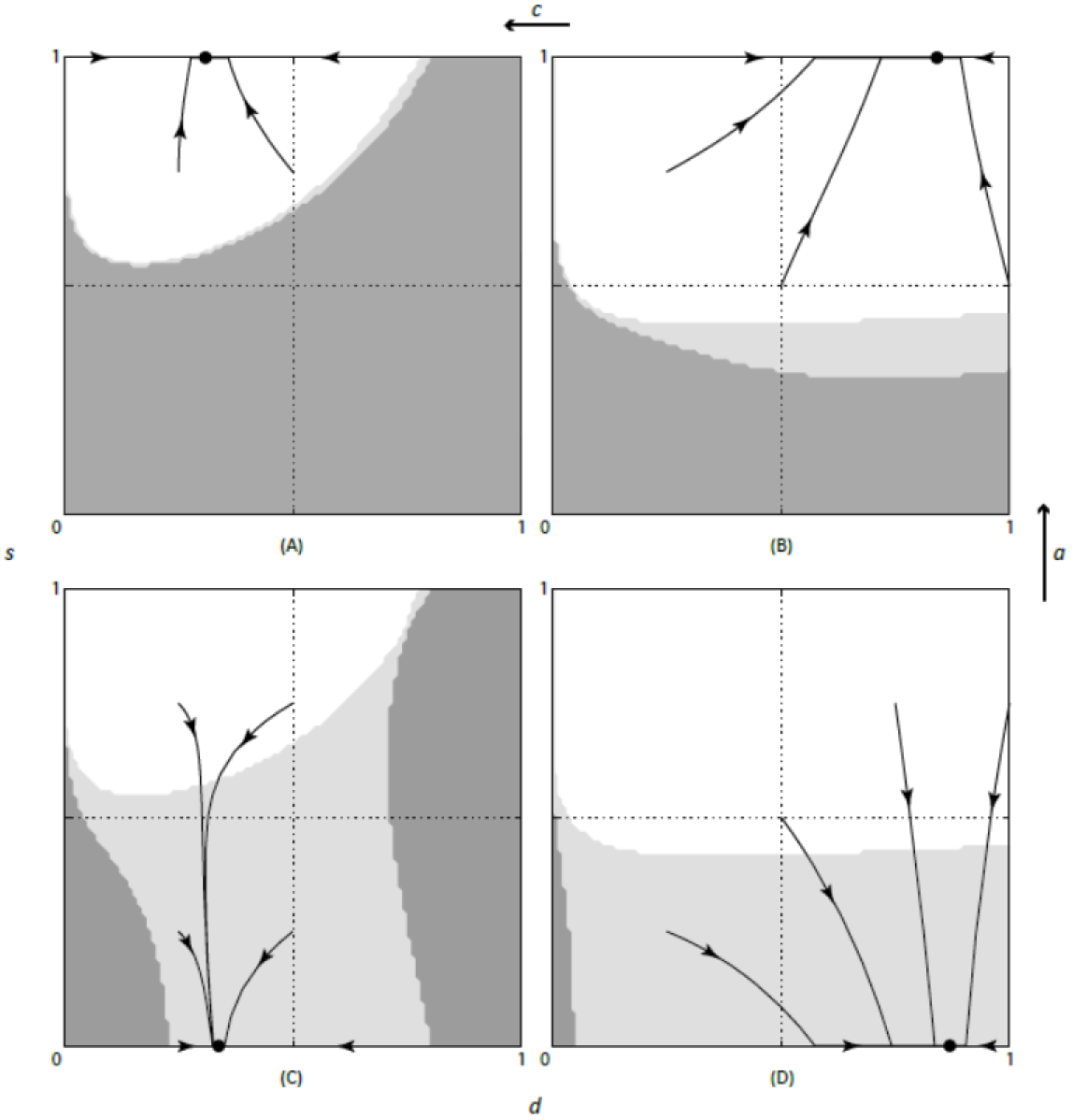
Consequences of cost of dispersal *c* and the Allee effect parameter *a* for evolution of dispersal rate *d* (horizontal axis) and selfing rate *s* (vertical axis) and for viability versus extinction of the metapopulation. Lines and arrows represent the path and direction of dispersal and selfing rate evolution from different starting values. Filled circles represent (globally stable and unique) evolutionarily stable strategies (ESS). Shading denotes ecological scenarios: white for the Unconditional Viability scenario, dark grey for the Unconditional Extinction scenario and light grey for the Conditional Viability scenario. Moving from right to left, cost of dispersal *c* increases (from 0.1 to 0.75), causing a decrease of dispersal rate (from about 0.84 to 0.3). Moving from bottom to top, the Allee effect parameter increases (from 10 to 100), causing an increase in selfing rate (from 0 to 1). Other parameter values are *δ* (inbreeding depression) = 0.5, *e* (local population extinction rate) = 0.1, *f* (fertility) = 10, *b* (competition parameter = 0.01).

**Fig.2.**
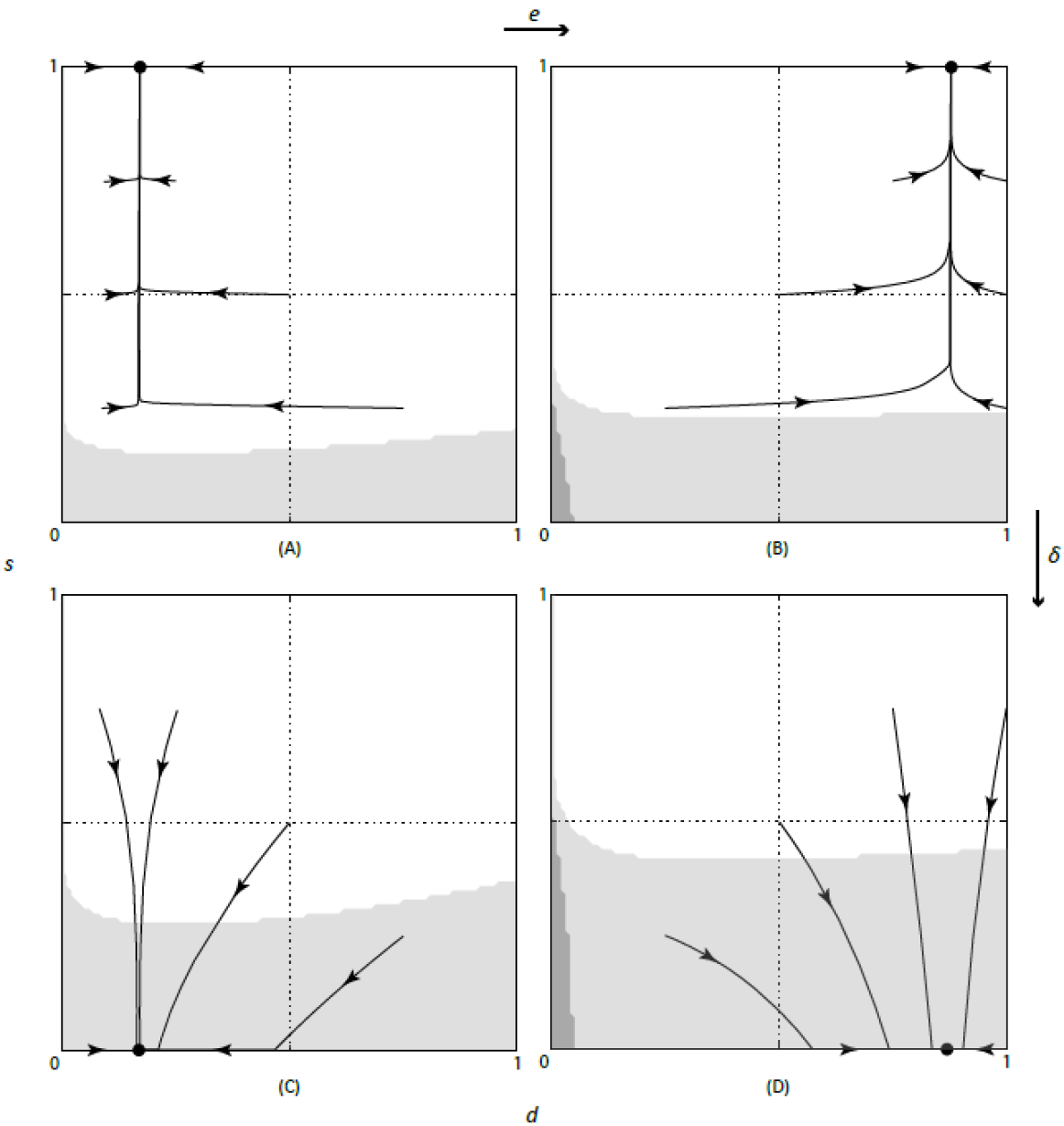
Consequences of inbreeding depression *δ* and local population extinction rate *e* evolution of dispersal rate *d* (horizontal axis) and selfing rate *s* (vertical axis) and for viability versus extinction of the metapopulation. Lines and arrows represent the path and direction of dispersal and selfing rate evolution from different starting values. Filled circles represent (globally stable and unique) evolutionary stable strategies (ESS). Shading denotes ecological scenarios: white for the Unconditional Viability scenario, dark grey for the Unconditional Extinction scenario and light grey for the Conditional Viability scenario. Moving from left to right, *e* increases (from 0.1 to 0.5), causing an increase of dispersal rate (from about 0.17 to 0.84). Moving from top to bottom, *δ* increases (from 0.1 to 0.5), causing a decrease in selfing rate (from 1 to 0). Other parameter values are *c* (cost of dispersal) = 0.1, *f*(fertility) = 10, *a* (Allee effect parameter) = 10, *b* (competition parameter = 0.01).

### 3.2 Sensitivity Analyses

The demographically structured HD model revealed a context dependent relationship between selfing and dispersal at ESS, with positive or negative relationships possible, depending which environmental or biological parameters varied (Fig. 3). Increasing local extinction rate increased both self-fertilisation and dispersal rates, producing a strong positive relationship between these traits and allowing for evolution of the GCS (Fig. 3a). Increasing the cost of dispersal increased selfing rate but decreased dispersal rate, producing a strong negative relationship (Fig. 3b). These are the only two parameters that produced marked joint evolution, as variation in other parameters affected selfing rate strongly but affected dispersal rate only weakly: higher selfing rates were associated with higher values of the competition parameter *b,* Allee effect parameter *a,* and lower values of fertility *f* and inbreeding depression *δ* (Fig. 3 c-d).

### 3.3 Selection on selfing and dispersal rates

Selection on selfing rate reflects the balance between the cost of inbreeding depression *δ* and the benefit of reproductive assurance, which depends on all the other parameters. Increasing the Allee effect parameter *a* (Fig. 3d) selects positively on selfing because this reduces cross-fertilisation of ovules/ova, and hence increases the benefit of selfing through reproductive assurance. By definition the Allee effect reduces cross-fertilisation more as local density becomes lower. Therefore, changing parameters so that density is reduced in some local populations should cause selection for higher selfing, and our results are consistent with this expectation. For instance, reducing fertility *f* increases the value of the ESS for selfing (Fig. 3d), which is expected because reducing propagule production will reduce local density, especially for younger (more recently recolonised) local populations. The increase in the ESS for selfing with increasing cost of dispersal *c* and increasing competition parameter *b* (Fig.3b, c), is likewise consistent with expectations based on the effect of these parameters on local density via propagule survival. The ESS for selfing also increases with increasing values of *e* (Fig. 3a), which is expected because increasing *e* leads to an increasing frequency of young, low-density local populations (i.e., a more left skewed distribution of local population density). In the approximated evaluation of fitness, the direction of selection on selfing depends only on *δ, a,* and *b,* and they have qualitatively the same effects as they do in the full HD model (Appendix B).

Selection on dispersal rate reflects the balance between a direct and negative effect of the cost of dispersal *c* (Fig. 3b) and the positive effect of escape from density-dependent resource competition. We thus attribute the increase in the ESS for dispersal under higher extinction rates (Fig. 3a) to an increased benefit of escape from competition when density is more heterogenous (Parvinen 2006). At low levels of *e*, most local populations are close to carrying capacity, so propagules will frequently disperse to a similar competition environment relative to the one in which they originate, resulting in selection for low dispersal rate. As *e* increases, the frequency of patches with low competition — i.e., empty patches, and low-density recently recolonised ones — increases. This increased heterogeneity in local density and local competition provides increased opportunity for propagules from older, high-density high-competition local populations to escape competition by dispersal, and hence the ESS for dispersal increases. However, heterogeneity in density and competition decreases again as *e* approaches its maximum, because then most populations have low density. This can lead to a decline in the ESS for dispersal although high *e* usually results in the metapopulation becoming extinct instead (results not shown, but see Parvinen 2006).

Evolution of selfing and dispersal differed from each other the HD model in that the ESS for selfing was almost always at extreme values (0 or 1) whereas that for dispersal never was. We attribute the result for dispersal to a negative eco-evolutionary feedback. Increasing dispersal rate reduces heterogeneity in local density of adult plants, reducing the benefit of escape from competition to be gained from any further increase in dispersal. Moreover, the higher the rate of dispersal, the greater the competition between dispersed propagules. Thus, as dispersal rate increases towards 1, the benefit of any further increase in dispersal becomes less until it is matches the cost of dispersal, causing dispersal rate to stabilises before it reaches 1. Similar logic explains why dispersal does not reach 0 either (Dornier et al. 2006). In contrast, although stable intermediate selfing rates 0 < *s* < 1 (mixed mating) are possible in this model (see Fig. 3 and Supplementary Materials), they only occur for very narrow ranges of parameter values. Most parsimoniously, selection of extremes may occur because there are thresholds in parameter values beyond which selection is always either positive or negative (Fisher 1941). This kind of effect occurs in our approximated analysis of selection, where thresholds separate parameter values producing positive versus negative selection on selfing (Appendix B). However, because selfing rate affects density, there could also be a positive eco-evolutionary feedback such that greater outcrossing success leads to generally higher local density and increased selection for outcrossing while reduced outcrossing success leads to generally lower local density and increased selection for selfing (Dornier et al. 2008).

**Fig. 3.**
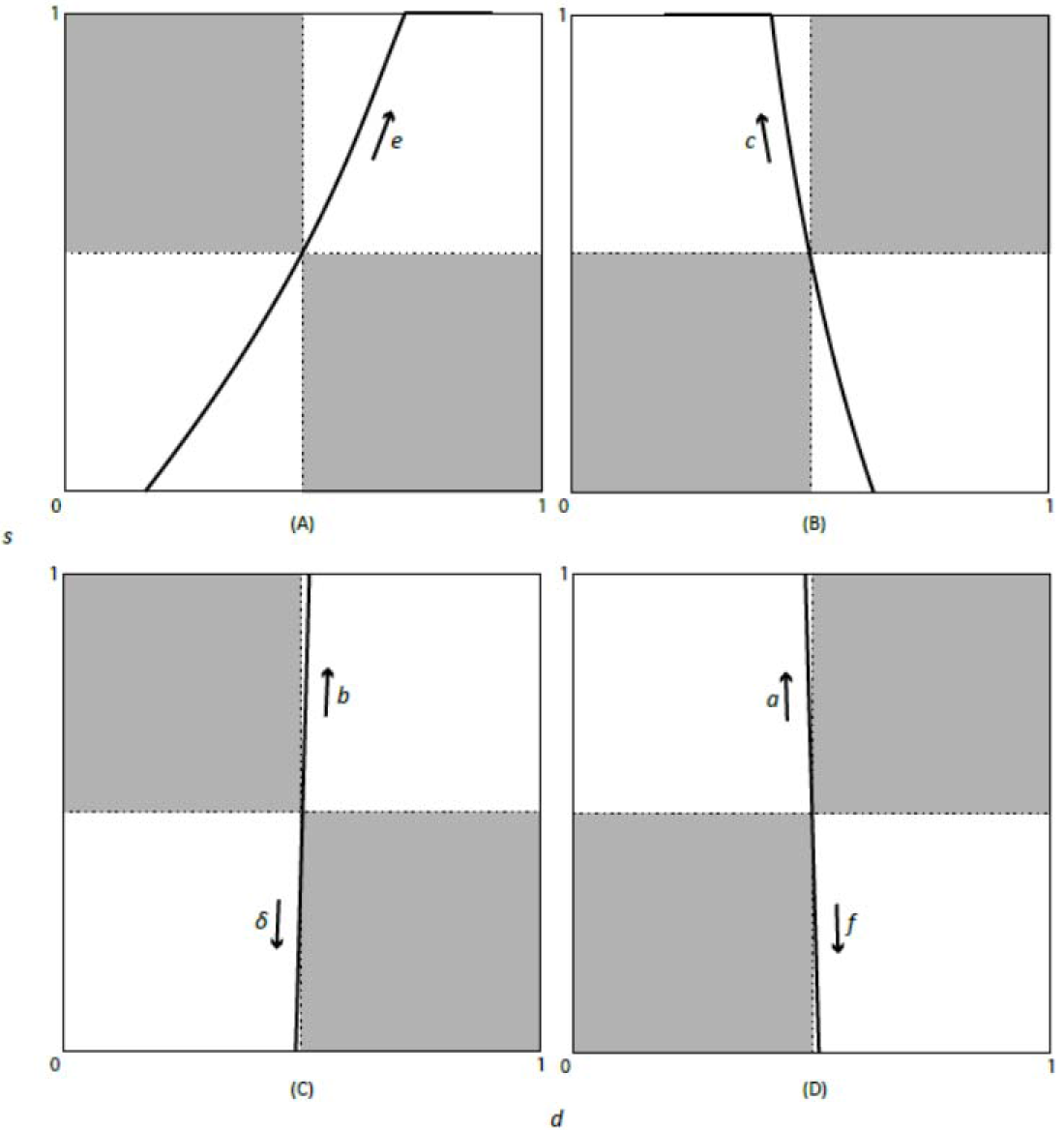
Sensitivity analysis showing how the dispersal-self fertilisation rate ESS shifts between quadrants of the D-S space. *e* = local extinction rate, *c* = cost of dispersal, *b* = competition parameter, *δ* = inbreeding depresson; *a* = Allee effect parameter and *f* = fecundity. Their effects on ESS are shown for *e* and *c* separately (top panels) and with remaining parameters grouped in pairs with opposite effects (bottom panels). Starting parameter values are *e* = 0.21, *c* = 0.1, *δ* = 0.10685, *b* = 0.01, *a* = 10, *f* = 10 and starting dispersal and selfing rate values are both 0.5. Parameter ranges explored are: *e:* 0.1−0.7; *c*: 0−0.8; *δ:* 0−1; *b:* 0.005−0.1; *a:* 0−100; *f*: 5−100. Shading of quadrants is not symbolic.

## 4. Discussion

### 4.1 Selection on selfing and dispersal rates

The HD model predicts a strong positive evolutionary relationship between selfing and dispersal when local extinction rate varies, offering a theoretical explanation for the positive relationship found in empirical studies (van Kleunen and Johnson 2007, Darling et al. 2008, De Waal et al. 2014, Grossenbacher et al. 2015). On the other hand, a strong negative relationship is predicted when cost of dispersal varies, and weak relationships are predicted when other parameters vary (Fig. 3). This context dependent relationship contrasts to the strictly negative one shown by the HP model (Cheptou and Massol 2009).

Varying local extinction causes a positive relationship between selfing and dispersal in the HD model (Fig. 3a) because as extinction rate increases, the frequency of local populations with low density increases in the metapopulation. This in turn favours higher selfing through the reproductive assurance benefit and higher dispersal through the escape from competition benefit, allowing evolution of the GCS of high dispersal and high selfing. In the HP model (Cheptou and Massol 2009), there is an identical (local) density of reproductive individuals in all patches. Therefore, propagule production of patches depends only on per-capita seed production and, as selfing rate increases under the influence of selection, propagule production of unpollinated patches approaches that of pollinated patches. This results in increasingly homogeneous competition conditions, reducing the benefit of dispersal. Hence the HP model predicts a strictly negative relationship between selfing and dispersal and no GCS (excepting a special case where selfed and outcrossed seeds evolve different rates of dispersal, Iritani and Cheptou 2017). In contrast, in the HD model, heterogeneity in seed production and competition arises not only from heterogeneity in per-capita reproduction, as a function of local density, but also directly from local density. Because extinction-recolonisation dynamics ensure heterogeneity in density, selfing does not homogenise seed production and competition in the HD model, and low dispersal does not have to be selected with high selfing.

The HD model predicts that selfing rate almost always evolves to the extreme values of 0 and 1, coarsely capturing the empirical distribution of selfing rates. The latter is bimodal with peaks close to 0 and 1, but with a substantial fraction of species with intermediate selfing (Moeller et al. 2017). Many previous models that also have limited complexity in selection on self-fertilisation predict only complete selfing or outcrossing (Lande and Schemske 1985; Cheptou 2004; Dornier et al. 2008). Models that predict evolutionarily stable mixed mating as a general result (reviewed in Goodwillie et al. 2005) include details that are absent from these models and the HD model, for instance allowing inbreeding depression to evolve or to fluctuate (e.g., Cheptou and Schoen, 2002) and, for plants, recognising that pollinators carrying out some self-pollination as well as cross-pollination (i.e., pollen discounting, Holsinger 1986, Porcher and Lande, 2005). Thus, the near absence of mixed mating in predictions of the HD model likely reflects the absence of these processes, which were not needed to meet the aims of this study.

### 4.2 Spatial and temporal variation in environmental pressures

Sensitivity analyses can be interpreted to shed light on how environmental gradients may result in stable or expanding range limits through ecological and evolutionary effects (Sagarin and Gaines 2002, Sun and Cheptou 2012, Hargreaves and Eckert 2014). Although a pattern of increased self-fertilisation ability at the range margin is common (e.g. Moeller 2006, Busch 2005, Barrett et al. 1989), quantification of both dispersal and selfing in relation to range position remains rare (Darling et al. 2008, de Waal et al. 2014). The HP model predicts range limits may be fixed (pinned) if pollinator failure increases towards the range edge, because this leads to an increase in selfing, which in turn leads to a decrease in dispersal (Sun and Cheptou 2012). In the HD model the same effect is produced by increasing dispersal cost towards the range edge (Fig. 3b), which could occur due to increasing distance between patches. In contrast, the HD model predicts that a gradient of local extinction rate from range centre to range edge would lead to a transition from outcrossing with low dispersal in the range centre to the GCS at the edge (Fig. 3a). Consistent with this prediction, in the dune plant *Abronia umbellata* both selfing and dispersal ability increase towards the range edge (Darling et al. 2008). The presence of local populations showing the GCS at the range edge should favour range expansion by long distance dispersal, as Baker envisioned for the colonisation of oceanic islands (Baker 1955).

The HD model also suggests how joint evolution of selfing and dispersal might proceed during spread of invasive species. Range expansion is likely to involve repeated colonisation events involving mate limitation Allee effects, accompanied by a decrease in habitat occupancy, and hence an increase in heterogeneity in competition, from the core to the invasion front (Simmons and Thomas, 2004). Increases in both self-fertilisation and dispersal rates could thus be favoured by conventional selection, due to mate limitation and escape from competition. However, they could also both be selected by spatial sorting over repeated colonisation events (Perkins et al. 2013, Ochocki and Miller 2016, Williams et al. 2016). Evolution of the GCS at invasion fronts thus seems plausible. Thus far, however, evolution of increased dispersal (Simmons and Thomas 2004, Phillips et al. 2006, 2008, 2010, Monty and Mahy 2010), but not selfing (Colautti et al. 2010) has been documented at the invasion front. However, the finding of higher selfing ability in the introduced range than the native range for several species of invasive plants (Petanidou et al. 2012) is consistent with this scenario.

### 4.3 Model limitations and future directions

Most simplifying assumptions made in our definition of fitness (section 2.1) should have quantitative rather than qualitative effects on our results. For instance, we omit the automatic transmission advantage of selfing (Fisher 1941) but doing so simply increases selection for selfing (Fig. S2). Allowing inbreeding depression to evolve should have the same effect, as lower inbreeding depression evolves at higher rates of selfing (Dornier et al. 2008, Massol and Cheptou 2011a). Conversely, allowing perenniality or a propagule bank (dormant propagules) could reduce selection for selfing by increasing local population density across the metapopulation (Pannell and Barret 1998). It would, however, be worthwhile to assess robustness of our findings to different patterns of spatiotemporal heterogeneity. For instance, local extinction probability could be allowed to increase with patch age, to represent successional environments (Comins, 1980), or heterogeneity in carrying capacity among patches could be introduced. In addition, as the HD model is very general, extensions tailored to reflect specifics of biology and parameterised realistically would allow more robust tests in particular taxonomic groups. Nevertheless, some important phenomena may not be compatible with the current modelling framework.

The HD model presented here does not account for higher relatedness of plants within than among local populations in the metapopulation, i.e., genetic structure. As this may moderate selection on both dispersal and selfing (Ravigné et al. 2006; Tonnabel et al. 2014; Welsford et al. 2016), investigating its effects on their joint evolution should be particularly rewarding. Cross-fertilisation between related individuals (biparental inbreeding) is frequent when there is genetic structure, and given that related individuals are more likely to share deleterious alleles than unrelated ones, this leads to inbreeding depression, albeit at a lower level than for selfing. This could makes it easier for selfing to evolve by reducing the difference in performance between offspring from selfing versus outcrossing (Lloyd 1979). Moreover, with genetic structure, dispersal has additional advantages for avoidance of (biparental) inbreeding and kin competition (Motro 1991, Perrin and Mazalov, 1999). These effects would most easily be dealt with using a spatially explicit modelling approach, as opposed to the spatially implicit approach used in this study.

### 4.4 Conclusions

The HD model shows that a positive relationship between dispersal and selfing is a likely outcome of evolution due to heterogeneity in local density, which promotes evolution of dispersal for escape from competition, and evolution of selfing for reproductive assurance against mate limitation. Consequently, the GCS can be selected under relatively high local extinction rates provided that inbreeding depression and cost of dispersal are not excessive. This contrasts to the HP model which predicts a strictly negative relationship between selfing and dispersal and no evolution of the GCS (Cheptou and Massol 2009, Massol and Cheptou 2011, Sun and Cheptou 2012). Thus by showing that both positive and negative relationships are possible, depending on context, our HD model extends our understanding of the joint evolution of these traits and reconciles theory with empirical findings (van Kleunen and Johnson, 2007; Darling et al. 2008; De Waal et al. 2014; Grossenbacher et al. 2015). Moreover, in considering selection on selfing from mate limitation, it provides a theoretical basis for understanding the joint evolution of selfing and dispersal in animals, whereas previous theory was only applicable to plants.

## Appendix

### A. Evaluation of mutant fitness

When a mutant individual appears in the metapopulation settled at its equilibrium, we can compute the mutant fitness *R*(*d, s, d', s'*) as the number of successfully established immigrant individuals that this mutant is expected to produce during the mutant population lifetime. If we assume that the mutant will appear at the beginning of the reproductive season in a patch of age *τ* ≥ 1 (a mutant individual cannot appear in a patch of age *τ* = 0 due to local extinction), that no new mutant immigrants are expected to arrive in the patch (the mutants being rare), that the mutant population and mutant seeds produced are always negligible compared to residents, and that no local extinctions will happen, the mutant population dynamics is described by

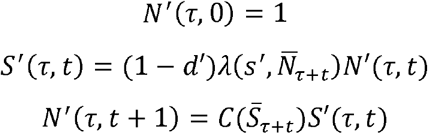

with *τ* ≥ 1. Note that, for the mutant reproduction, we are using the same function *λ* introduced for the resident, simply evaluating it with the mutant trait *s'*. Estimation of fitness in the HD model is simplified in that it ignores the transmission advantage: i.e., a propagule arising from self-fertilisation contains two copies of the maternal parent’s genes whereas a propagule from cross-fertilisation contains only one (Fisher 1941). This means that we underestimate selection on self-fertilisation, so our assessment of conditions favouring evolution of self-fertilisation, and hence the GCS, is conservative in this respect. Although we could have used the function of Dornier et al. 2008 (Equation 10) which incorporates the transmission advantage, this equation is still simplified and overestimates selection on self-fertilisation compared to an accurate sexual model (Parvinen and Metz 2008). Nevertheless, simulations on a version of the HD model incorporating the transmission advantage sensu Dornier et al. 2008 confirms that omitting it does not qualitatively affect the results (Supplementary Material).

The number of emigrant seeds that this mutant is expected to produce during the entire lifetime of the patch (before it goes extinct) is thus given by

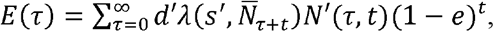

with (1 – *e*)*^t^* the probability that local extinction has not happened in *t* time steps.

We must now account for the age distribution *p_τ_* of patches where the mutant can appear. Moreover, to become an immigrant mutant individual, emigrant mutant seeds must survive dispersal and competition. Thus, mutant invasion fitness is given by

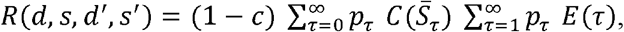

where (1 – *c*) is the probability of mutant emigrant seeds to survive dispersal, the first sum is the expected probability of immigrant mutant seeds to survive competition and become immigrant mutant individuals, and the second sum is the expected number of emigrant mutant seeds produced by the mutant population during its lifetime.

Following the approach of Parvinen 2006 (see Appendix A in particular), the number of emigrant seeds can be computed recursively, i.e., the per-capita number of emigrants produced by a mutant appearing in a patch of age *τ* are given by two contributions: the first is the number of seeds directly dispersed by that mutant, while the second represents the seeds that stay in the patch. If they survive competition and local extinction, they can be considered as mutants appearing in a patch of age *τ* + 1, that in turn will produce emigrant seeds. In formulas,

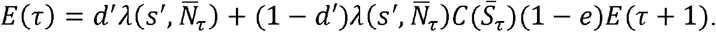

If we assume that the population dynamics has converged to its attractor, we can practically set an upper limit L for the age *τ* of the patches so that

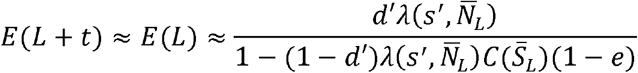

for each *t >* 0. Therefore we can set *R*(*L*) = *E*(*L*)(1 – *e*)*^L^* (where (1 – *e*)*^L^* is the probability that a randomly chosen patch has age L or larger) and recursively compute, for *t* = 1, ..., *L* – 1,

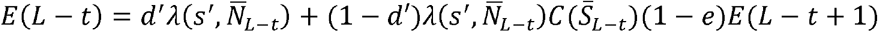

and

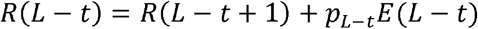

until obtaining *R* (1), that approximates the expected number of emigrant mutant seeds produced by a mutant population during its lifetime (recall that reproductive mutants cannot appear in patches of age *τ* = 0, so that *E*(0) = 0 and *R*(0) = *R*(1)). The mutant fitness will then be approximated by

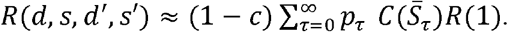

The selection gradient used for the evolutionary simulations is the derivative of the invasion fitness with respect to the mutant traits, i.e.,

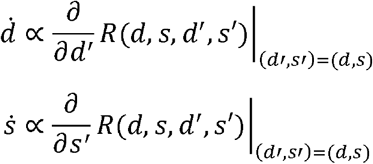

### B. Approximated evaluation of fitness

To gain insight into selection on dispersal and selfing in the HD model, we perform an approximated analysis of the selection gradients. We do this by assuming that all the non-extinct patches have reached the stationary maximum local population 1/*b*, in other words, neglecting distribution of local population density. This approximation is thus most helpful for explaining results of the full model that do not depend on the frequency distribution of local density (see Discussion).

We approximate *R* (1) = *R*(*L*) = *p_L_E*(*L*) to find approximated expressions for the sign of the selection gradients on *d* and *s* and thus of the trait values at ESS, *d_ESS_* and *s_ESS_*. Taking the derivatives of *E*(*L*) with respect to the mutant traits *d'* and *s'*, the sign of the selection gradient on selfing is given by the sign of (1 – *δ*) – 1/(1 + *ab*), while the sign of the selection gradient on dispersal is given by the sign of 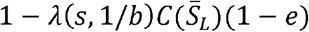, with 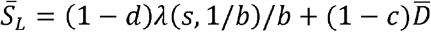 and 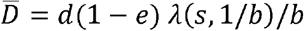.

Notably, the expression for selfing does not depend on the traits (*d, s*) themselves, supporting our observation of selection for extreme mating strategies (either full selfing or full outcrossing) in the HD model. On the other hand, direction of selection on dispersal is given by a balance between extinction risk, fertility, and survival of competition, i.e., the sign of 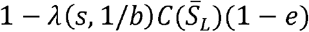, where 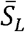 is the number of seeds in the non-extinct patches that has to survive competition through function *C*(*S*) = 1/(1 + *bs*). The former number of seeds is itself a function of dispersal *d*, fertility *λ*(*s*, 1/*b*), local population size in the oldest patches 1/*b*, cost of dispersal *c*, and extinction rate *e*. Notice that instead this expression depends on the traits (*d, s*), thus it can be used to find approximated implicit expressions for the two traits at the ESS by solving for neutral selection, obtaining (by also substituting the expression for *λ*, *C*(*s*), and *D*, see main text and Appendix A)

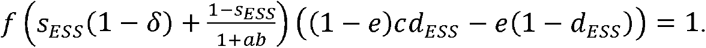

This expression can be used to assess and estimate the positive vs. negative (*d, s*) relationship. In fact, perturbing (taking the derivative with respect to) any parameter one can find an implicit expression of the resulting perturbations on the ESS. For example, perturbing the extinction rate *e* would lead to (remember that *d_ESS_* and *s_ESS_* are themselves implicit functions of all parameters)

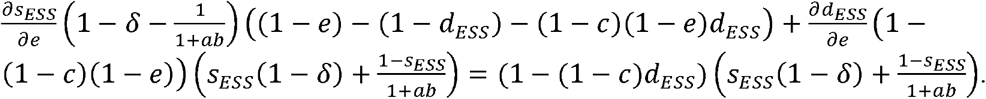

Such expressions can be used to estimate for a generic parameter *p* as a positive 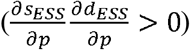 vs. negative 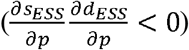 (*d,s*)-relationship for perturbations in any
parameter *p*.

However, *s_ESS_* is generically either 0 or 1, depending on the sign of (1 – *δ*) – 1/(1 + *ab*). We can exploit this by obtain a simplified expression for *d_ESS_* for each of these two cases.

**Table B1:** Selfing and dispersal rate at ESS in the simplified model.

| | $s_{ESS}$ | $d_{ESS}$ |
| --- | --- | --- |
| $(1 - \delta) < 1/(1 + ab)$ | 0 | $\frac{1 + ef(1 - \delta)}{f(1 - \delta)(e + c(1 - e))}$ |
| $(1 - \delta) > 1/(1 + ab)$ | 1 | $\frac{1 + ab + ef}{f(e + c(1 - e))}$ |

Thus, the effect of each parameter can be computed. As for selection on selfing, *δ* (respectively, *a* and *b*) have a negative (respectively, positive) effect. If the selection gradient is close to singular ((1 – *δ*) = 1/(1 + *ab*)), then a small parameter perturbation can trigger the transition of *s_ESS_* from 0 to 1 or vice versa. The sign of the effect of each parameter on dispersal is given in the following table.

**Table B2:** Effect of parameter perturbations on the dispersal rate at ESS in the simplified model.

| Parameter | $\frac{\partial d_{ESS}}{\partial p} (s_{ESS} = 0)$ | $\frac{\partial d_{ESS}}{\partial p} (s_{ESS} = 1)$ |
| --- | --- | --- |
| $e$ | $\frac{cf - (1 + ab)(1 - c)}{f(e + c(1 - e))^2}$ | $\frac{cf(1 - \delta) - (1 - c)}{f(1 - \delta)(e + c(1 - e))^2}$ |
| $c$ | $-\frac{(1 - e)(1 + ab + ef)}{f(e + c(1 - e))^2}$ | $-\frac{(1 - e)(1 + ef(1 - \delta))}{f(1 - \delta)(e + c(1 - e))^2}$ |
| $f$ | $-\frac{(1 + ab)}{f^2(e + c(1 - e))}$ | $-\frac{1}{f^2(1 - \delta)(e + c(1 - e))}$ |
| $\delta$ | 0 | $\frac{1}{f(1 - \delta)^2(e + c(1 - e))}$ |
| $a$ | $\frac{b}{f(e + c(1 - e))}$ | 0 |
| $b$ | $\frac{a}{f(e + c(1 - e))}$ | 0 |

Table B2 can then be simplified to table B3

**Table B3:** Qualitative effect of parameter perturbations on the dispersal rate at ESS in the simplified model.

| Parameter | $\frac{\partial d_{ESS}}{\partial p} (s_{ESS} = 0)$ | $\frac{\partial d_{ESS}}{\partial p} (s_{ESS} = 1)$ |
| --- | --- | --- |
| $e$ | $cf - (1 + ab)(1 - c)$ | $cf(1 - \delta) - (1 - c)$ |
| $c$ | <0 | <0 |
| $f$ | <0 | <0 |
| $\delta$ | 0 | >0 |
| $a$ | >0 | 0 |
| $b$ | >0 | 0 |

